# Acute stress induces circuit-specific alterations in mesolimbic and nigrostriatal inhibitory transmission and potentiates operant learning

**DOI:** 10.64898/2026.06.04.729088

**Authors:** Helena de Carvalho Schuch, Joyce Woo, Hannah Kugler, Kelly Jiang, Alexey Ostroumov

**Author notes:** Address all correspondence to Alexey Ostroumov, Department of Pharmacology and Physiology, Georgetown University, Washington, DC 20007, USA.

## Abstract

Acute stress can facilitate learning about stimuli that predict rewarding outcomes, yet whether stress similarly potentiates acquisition of reward-directed actions remains less well understood. Reward learning broadly depends on dopamine signaling within mesolimbic and nigrostriatal pathways. Dopamine transmission in the mesolimbic system is traditionally associated with cue-reward learning, whereas nigrostriatal dopamine signaling has been implicated in movement vigor and the acquisition of instrumental actions. Stress can alter dopamine transmission within these circuits and thereby influence reward learning. Previous work demonstrated that acute stress downregulates the potassium-chloride cotransporter KCC2 in ventral tegmental area GABA neurons. These stress-induced adaptations in inhibitory transmission enhance mesolimbic dopamine signaling and potentiate associative learning. Here, we show that prior exposure to acute restraint stress facilitates acquisition of operant sucrose self-administration in male and female rats. Enhanced learning was associated with increased temporal coincidence of GABA release events onto dopamine neurons and increased excitability of GABAergic inputs, alterations previously linked to enhanced dopamine signaling. These stress-induced adaptations exhibited marked circuit specificity within mesolimbic and nigrostriatal systems, selectively affecting inhibitory transmission onto dopamine neurons projecting to the nucleus accumbens lateral shell and dorsomedial striatum. Importantly, pharmacological enhancement of KCC2 function with CLP290 normalized inhibitory transmission within these pathways and attenuated stress-induced potentiation of operant learning. Together, these findings identify circuit-specific alterations in midbrain inhibitory signaling induced by acute stress that contribute to enhanced reward learning.

## 1. Introduction

Acute stress profoundly influences reward-related behavior, which is essential for survival and well-being. In adaptive contexts, stress can facilitate learning about stimuli and actions that predict beneficial outcomes such as food, water, and reproduction. However, stress-induced alterations in reward learning can also contribute to the development of neuropsychiatric disorders, including post-traumatic stress disorder, substance use disorder, and major depressive disorder. Existing evidence indicates that acute stress enhances classical conditioning, in which environmental cues acquire predictive value for rewarding outcomes (Stelly et al., 2020). In contrast, the impact of stress on operant reward learning, in which organisms learn the specific actions that lead to reward delivery, is less well understood (Schettino et al., 2024). While previous studies have shown that stress can potentiate operant learning for addictive drug rewards (Carter et al., 2020; Garcia-Keller et al., 2016), it remains unclear whether acute stress similarly facilitates operant learning for natural rewards.

Stress-induced changes in operant learning likely involve modulation of mesolimbic and nigrostriatal dopamine (DA) systems in the brain (Wise, 2004). The mesolimbic DA pathway between the ventral tegmental area (VTA) and nucleus accumbens (NAc) primarily supports reinforcement and motivational processes that drive the initiation and persistence of reward-seeking actions (Ikemoto and Panksepp, 1999; Salamone and Correa, 2012). In parallel, nigrostriatal DA pathways between the substantia nigra pars compacta (SNc) and dorsal striatum are more directly involved in encoding action-outcome contingencies and selecting appropriate actions that support goal-directed behavior and its transition to habitual responding (Balleine and O’Doherty, 2010; Yin and Knowlton, 2006). Accumulating evidence suggests that DA neuron populations within the mesolimbic and nigrostriatal systems are further subdivided into anatomically and functionally distinct projections targeting specific subregions of the NAc and dorsal striatum, where they regulate different aspects of reward processing (De Jong et al., 2019; Engel et al., 2024; Lerner et al., 2015; Saunders et al., 2018). Stress may differentially modulate these projection-defined circuits through pathway-specific changes in synaptic input. However, the molecular and neuronal mechanisms underlying these stress-induced adaptations, and their contribution to reward learning, remain largely unknown.

We recently found that acquisition of reward learning depends on transient downregulation of the chloride transporter KCC2 in VTA GABA neurons, which are major inhibitory regulators of VTA DA neurons (Woo et al., 2025). Impaired chloride homeostasis synchronizes VTA GABA neuron activity, increasing coincident inhibitory input onto VTA DA neurons. This synchronized inhibitory input enhances phasic DA activity and DA release within lateral mesolimbic pathways, thereby facilitating learning of associations between stimuli and rewards (Woo et al., 2025). Because stress has also been shown to downregulate KCC2 and to facilitate operant responding for drugs of abuse (Ostroumov et al., 2016), we hypothesize that stress-induced KCC2 downregulation may also facilitate operant learning for natural rewards. Given the complementary roles of mesolimbic and nigrostriatal DA systems in operant learning, we further predict that stress-induced KCC2 downregulation differentially impacts inhibitory inputs onto DA neurons projecting to distinct downstream pathways across both the VTA and SNc.

To examine the interaction between stress and operant reward learning, we exposed rats to acute restraint stress and subsequently measured sucrose self-administration. Along with enhanced acquisition of sucrose self-administration, stress synchronized GABAergic input onto VTA and SNc DA neurons in a pathway-specific manner. Importantly, stress-induced increases in operant responding and coordinated inhibitory input onto DA neurons were reversed by enhancing KCC2 function. Together, these findings demonstrate that acute stress differentially alters GABAergic regulation of both mesolimbic and nigrostriatal DA pathways, contributing to enhanced operant reward learning.

## 2. Materials and methods

### 2.1 Animal Model

Wild-type and TH-Cre Long–Evans rats (250-400 g; both sexes; Harlan/Envigo/Inotiv) were housed in same-sex pairs under controlled temperature and humidity conditions on a 12 h light/dark cycle, with ad libitum access to food and water throughout the study. All behavioral experiments were conducted after a minimum of 5 days of handling the rats. No health or weight difference was detected between control and stressed animals at any point during the experimental manipulation. All procedures were conducted in accordance with the guidelines from the Institutional Animal Care and Use Committee (IACUC) at Georgetown University’s Medical Center.

### 2.2 Acute restraint stress

Broome-style rodent restrainer tubes (Plas Labs) were used to restrain rats for one hour. Behavioral responses to stress included increased vocalization, dander release, defecation, and urination (DeTurck and Vogel, 1982; Ostroumov et al., 2016). After stress, animals were rehoused with their non-stressed cage mate. All restraint stress tests were conducted during the initial hours of the dark light cycle (6:00-9:00 pm) and 15-20 hours prior to patch-clamp recordings or the start of behavioral experiments. The 15-hour interval between stress exposure and testing was selected to assess the long-term effects of acute stress on reward learning while allowing animals to recover from any transient physical discomfort associated with restraint.

### 2.3 Sucrose self-administration behavior

Standard operant conditioning chambers (Med Associates Inc.) were used for all self-administration experiments. Behaviorally naïve control and stressed rats were trained on a lever-pressing task under a fixed ratio 1 (FR1) schedule of reinforcement. Each active lever press resulted in delivery of a sucrose pellet paired with illumination of a cue light located directly above the lever. An inactive lever was also present during each session but had no programmed consequence. At the start of each session, the house light was illuminated, and a single sucrose pellet and cue light were delivered. Rats underwent daily 1-hour training sessions for 6 consecutive days. Prior to behavioral analysis, animals were excluded if they left two or more uneaten pellets on more than one day or exhibited indiscriminate lever pressing (>20 inactive lever presses in a session) during any session after the initial training session (Garcia-Keller et al., 2016). Acquisition of operant behavior was defined as the first of two consecutive sessions in which animals earned ≥10 rewards after correction for inactive lever pressing (active - inactive lever presses). Rats that met this acquisition criterion within the first six self-administration sessions received an additional four FR1 sessions, during which they continued to earn >10 rewards per session.

### 2.4 Intraperitoneal injections

For systemic administration, CLP290 was first dissolved in 40% β-cyclodextrin (20 mg/ml) and then diluted in saline to a final concentration of 10 mg/ml in 20% β-cyclodextrin (Gagnon et al., 2013; Ostroumov et al., 2020). The solution was adjusted to pH 5-6 using 10 N NaOH. CLP290 (10 mg/kg, i.p.) or vehicle (20% β-cyclodextrin) was administered 45 min prior to sucrose self-administration sessions on three non-consecutive days (Ostroumov et al., 2020; Thomas et al., 2018). To minimize stress, rats were habituated to brief hand restraint, and injections were performed in the home cage within the animal facility.

### 2.5 Stereotaxic surgeries

DA neurons projecting to different subregions of the NAc and dorsal striatum were labeled in TH-Cre rats using adeno-associated viruses delivered via stereotaxic surgery. Surgeries were performed using a stereotaxic apparatus (Stoelting Co.) under isoflurane anesthesia (2–3%), with body temperature maintained using an isothermal heating pad (Braintree). For labeling projection-defined DA neurons, AAV2/5-pCAG-Flex-eGFP (Addgene, Catalog # 51502-AAV5) was bilaterally injected into the NAc medial shell (AP +1.5, ML ±0.7, DV −7.4 to −7.5 mm), NAc core (AP +1.7, ML ±1.6, DV −6.6 to −6.7 mm), NAc lateral shell (AP +1.1, ML ±2.6, DV −7.9 to −8.0 mm), dorsomedial striatum (DMS; AP +0.25, ML ±2.3, DV −5.1 to −5.2 mm), or dorsolateral striatum (DLS; AP +0.25, ML ±3.8, DV −5.4 to −5.5 mm). Viruses were allowed to express for 3 weeks to enable labeling of midbrain DA neuron somata.

### 2.6 Patch-Clamp Electrophysiology

Rats were deeply anesthetized and brains were rapidly extracted for preparation of acute slices used in patch-clamp recordings. Horizontal midbrain slices (220 μm) containing the VTA and SNc were prepared using a vibratome (Leica Microsystems) in ice-cold oxygenated (95% O_2_/5% CO_2_) high-sucrose artificial cerebrospinal fluid (ACSF) containing (in mM): 205 sucrose, 2.5 KCl, 21.4 NaHCO_3_, 1.2 NaH_2_PO_4_, 0.5 CaCl_2_, 7.5 MgCl_2_, and 11.1 dextrose. After cutting, slices were immediately transferred to oxygenated normal ACSF containing (in mM): 120 NaCl, 3.3 KCl, 25 NaHCO3, 1.2 NaH_2_PO_4_, 2 CaCl_2_, 1 MgCl_2_, 10 dextrose, and 20 sucrose. Slices recovered in this solution for 40 min at 32°C and were subsequently equilibrated at room temperature for an additional hour. For experiments examining KCC2 enhancement, slices were incubated in ACSF containing CLP290 (10 μM) for an additional hour prior to recording.

For electrophysiological recordings, slices were transferred to a recording chamber (Scientifica) and continuously perfused with oxygenated ACSF (2-3 ml/min, 32°C). Recording electrodes were made of thin-walled borosilicate glass (1.12 mm inner diameter, 1.5 mm outer diameter; World Precision Instruments) and had resistances of 1-1.5 MΩ when filled with a high-chloride internal solution containing (in mM): 135 KCl, 12 NaCl, 2 Mg-ATP, 0.5 EGTA, 10 HEPES, and 0.3 Tris-GTP (pH 7.2-7.3). Projection-defined DA neurons were identified by Cre-dependent eGFP fluorescence using epifluorescence illumination (470 nm, CoolLED, Olympus). GABAergic input onto VTA and SNc DA neurons was assessed by recording spontaneous inhibitory postsynaptic currents (sIPSCs) in whole-cell voltage-clamp configuration at a holding potential of −60 mV. Ionotropic glutamatergic transmission was blocked by bath application of AP5 (20 μM, HelloBio) and DNQX (50 μM, HelloBio). GABA_A_ receptors on midbrain GABA neurons were additionally stimulated with bath-applied diazepam (Sigma). Signals were acquired using a Multiclamp 700B amplifier, filtered at 10 kHz, and digitized at 20 kHz using a Digidata 1550B interface and pClamp 11 software (Molecular Devices). Data were analyzed offline using Clampfit 11 (Molecular Devices).

### 2.7 Immunohistochemistry and microscopy

To verify viral labeling of DA neurons and the anatomical localization of labeled neurons within the VTA and SNc, rats were deeply anesthetized and transcardially perfused with 50 mL phosphate-buffered saline (PBS; EMD Millipore), followed by 50 mL 4% paraformaldehyde (PFA, Santa Cruz Biotechnology). Brains were post-fixed in 4% PFA and cryoprotected in ascending sucrose solutions prepared in PBS (10%, 20%, and 30%). Tissue was subsequently embedded in OCT and stored at −80°C prior to sectioning. Coronal brain sections (30 μm) containing the VTA and SNc were collected as free-floating sections in PBS. Sections were incubated in blocking solution containing 3% normal goat serum (NGS) and 0.3% Triton X-100 (Sigma) in PBS, followed by incubation with rabbit anti-tyrosine hydroxylase (TH) primary antibody (1:1000; Invitrogen) for 12-16 hours at 4°C. Sections were then incubated with goat anti-rabbit Alexa Fluor 647 secondary antibody (1:500; Invitrogen) for 3 hours at room temperature. Following staining, sections were mounted onto microscope slides using Vectashield mounting medium (Vector Laboratories) and stored at −20°C until imaging (Leica Microsystems).

### 2.8 Anatomical confirmation of injection sites

To verify AAV injection sites within targeted accumbal and striatal subregions, coronal brain blocks containing the injection sites were collected during preparation of slices for patch-clamp experiments and post-fixed in 4% PFA for at least 24 h. Tissue sections (150 μm) containing the striatal regions of interest were subsequently prepared using a vibratome (Lancer), and injection sites were verified by light microscopy (Olympus).

### 2.9 Statistical Analysis

Data are presented as mean ± SEM, and statistical significance was defined as p < 0.05. Repeated measures ANOVA (RM ANOVA) was used to analyze active and inactive lever presses during sucrose self-administration. Two-tailed unpaired t-tests were used to compare mean differences between active and inactive lever presses, as well as between groups for sIPSC frequency, synchrony, and amplitude. For analysis of diazepam-induced changes in sIPSCs, raw data (Hz) were converted to a percentage of baseline, with the last 30-60 seconds before diazepam application used as the baseline period. All statistical analyses were performed using Prism (GraphPad).

## 3. Results

### 3.1 Stress potentiates the acquisition of sucrose self-administration

We first examined how a single episode of restraint stress influences subsequent learning of a reinforcing behavior using operant sucrose self-administration. Male and female rats were exposed to restraint stress (1 hour) approximately 15 hours prior to the first self-administration session (Figure 1A). This interval was selected to assess lasting effects on neural circuits rather than the immediate effects of the stressor (Ostroumov et al., 2016). Pre-exposure to stress increased operant responding for sucrose across first six daily sessions compared to non-stressed controls (Figure 1B). Inactive lever responding did not differ between groups (Figure 1C). Consequently, the difference between active and inactive lever presses was significantly greater in stressed animals than in controls (Figure 1D, control: 6.7 ± 2.8, stress: 19.2 ± 4.5). Using a predefined acquisition criterion of ≥10 presses on the active versus inactive lever for at least two consecutive days, 57.9% of stressed animals acquired sucrose self-administration within the first six sessions, compared to 31.3% of control animals (Figure 1D). Among animals that met the acquisition criterion during the first six sessions, stressed and control rats exhibited similar levels of responding during days 7-10 (Figures 1E-G). Together, these results suggest that prior stress potentiates acquisition but not the expression of sucrose self-administration.

**Figure 1.**
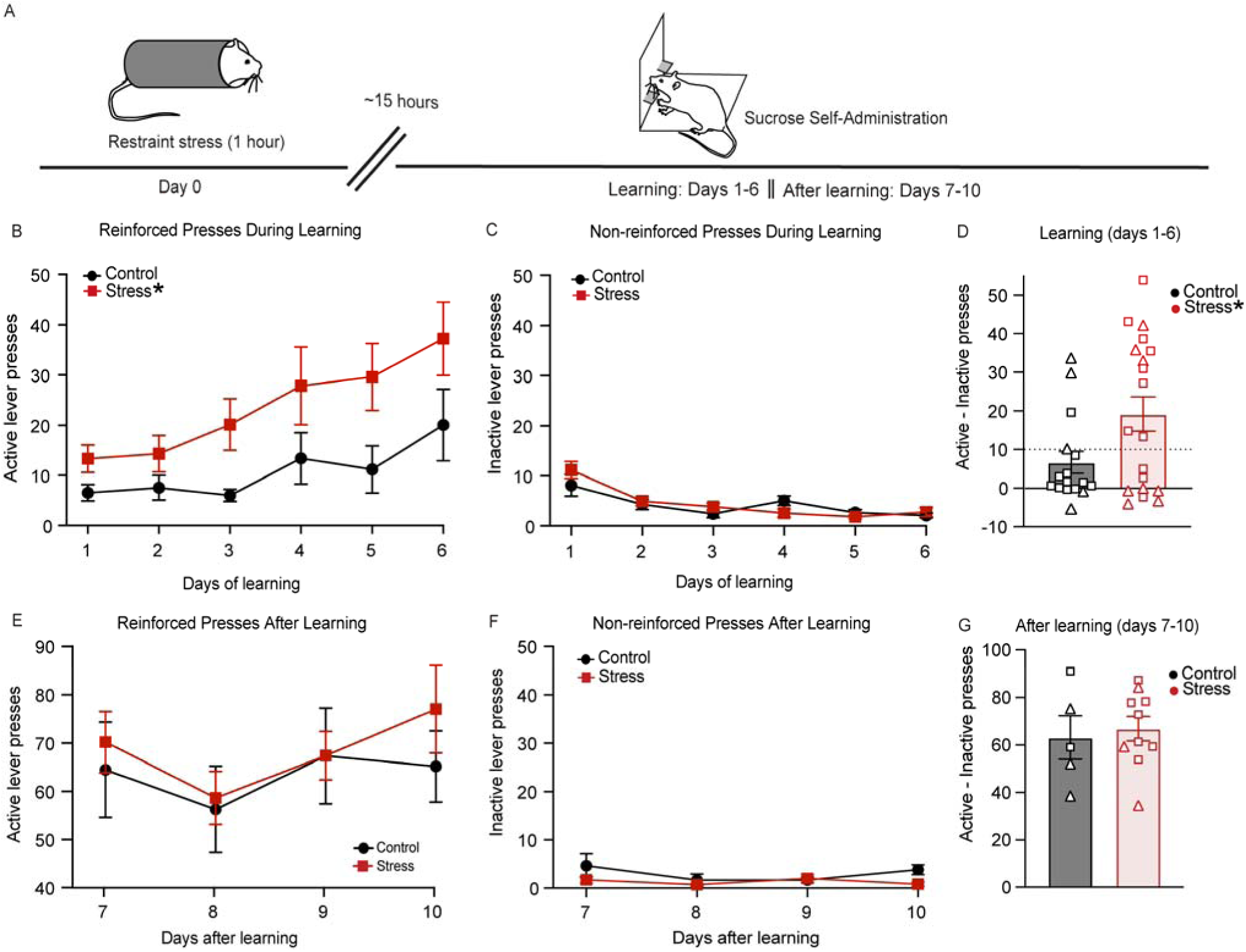
Stress potentiates the acquisition of sucrose self-administration. A) Rats were subjected to a single 1-hour restraint stress approximately 15 hours before the first sucrose self-administration session. Active and inactive lever presses were measured in control and stressed rats during the first 6 days of training, and for an additional 4 days in animals that acquired sucrose self-administration. B) Stressed rats exhibited increased active lever pressing during the first 6 days of sucrose self-administration compared to controls (*p = 0.0272, RM ANOVA, treatment: F (1, 33) = 5.345, n = 19 stress, 16 control rats). C) Inactive lever presses did not differ between stressed and control groups (p = 0.6239, RM ANOVA). D) The mean daily difference between active and inactive lever presses was greater in stressed animals compared to controls (p = 0.0297, two-sided t-test, n = 16, 19 rats). Triangles represent females and squares represent males. The dashed line indicates the acquisition criterion (≥10 presses on the active versus inactive lever). E) In animals that acquired sucrose self-administration within the first 6 sessions, active lever pressing did not differ between stressed and control groups (p = 0.5824, RM ANOVA, n = 10, 6 rats). F) Inactive lever presses during sessions 7–10 did not differ between stressed and control animals that acquired sucrose self-administration (p = 0.1429, RM ANOVA). G) The mean daily difference between active and inactive lever presses did not differ between stressed and control animals that acquired sucrose self-administration (p = 0.7096, two-sided t-test). Triangles represent females and squares represent males.

### 3.2 Stress alters GABAergic input onto NAc lateral shell-projecting DA neurons

Stress can potentiate reward-related behaviors by impairing KCC2 function in VTA GABA neurons, a mechanism previously shown to enhance Pavlovian learning through altered inhibitory control over DA projections to the NAc lateral shell (Ostroumov et al., 2016; Woo et al., 2025). To test whether acute stress impairs KCC2 function in this pathway, we retrogradely labeled DA neurons by injecting male and female TH-Cre rats with a Cre-dependent AAV expressing GFP into the NAc lateral shell (Figures 2A-B). Following 1 hour of restraint stress, we assessed stress-induced changes in GABA release by measuring the frequency of spontaneous inhibitory postsynaptic currents (sIPSCs) in GFP-positive VTA DA neurons 15 hours later (Figure 2C). KCC2 downregulation in VTA GABA neurons has previously been associated with increased synchronization of spontaneous GABAergic input onto DA neurons (Moore et al., 2018; Woo et al., 2025). Consistent with this, we observed an increase in the temporal clustering of GABA release events (Figure 2D, top black and red traces with insets). Quantitative analysis revealed that after stress, sIPSC frequency in lateral shell-projecting DA neurons remained unchanged (control: 4.3 Hz ± 0.8, stress: 4.6 Hz ± 0.7, p = 0.7566, n = 6, 9 cells in 3, 5 rats), but there was a significantly higher percentage of sIPSCs with interevent intervals <10 ms compared to non-stressed controls (Figure 2E, 11.3 ± 2% in stressed animals versus 2.5 ± 0.8% in control animals). To confirm the role of KCC2 downregulation, we recorded sIPSCs in slices treated with the KCC2 activator CLP290 (10 µM, 1 h), which decreased the percentage of sIPSCs with interevent intervals <10 ms to control levels (Figure 2E): 2.8 ± 0.7% in stress versus 1.4 ± 0.7% in control animals. Importantly, CLP290 did not affect the overall sIPSC frequency (3.2 ± 0.5 Hz in control and 4.7 ± 0.8 Hz in stressed rats, p = 0.1149), suggesting that stress-induced KCC2 downregulation enhanced synchronized GABA release onto DA neurons without altering total GABAergic input. Our previous study showed that increased GABAergic synchrony potentiates phasic DA firing and DA release in the NAc lateral shell (Woo et al., 2025). To further confirm the impact of KCC2 modulation on GABA release onto DA neurons, we used a pharmacological approach. KCC2 downregulation leads to VTA GABA neuron excitation during intense GABA_A_ receptor activation, and GABA_A_ receptors in VTA GABA neurons are more sensitive to the benzodiazepine diazepam than in DA neurons (Tan et al., 2011, 2010). As expected, bath application of diazepam (5 µM) increased sIPSC frequency in DA neurons from stressed animals compared to controls (Figure 2F): 132.7 ± 11.4% of basal frequency in stressed animals versus 74.9 ± 4.4% in controls. This effect was blocked by pre-incubation with CLP290 (Figure 2F; 64.9 ± 4.7% in stressed animals versus 73 ± 5.4% in controls). Importantly, diazepam did not alter sIPSC amplitude between stressed and control animals, indicating that the increased sIPSC frequency was due to enhanced presynaptic GABA release, rather than postsynaptic adaptations (control = 101.6 ± 8.1% of basal; stress = 104.8 ± 15.7% of basal frequency, p = 0.8411).

**Figure 2.**
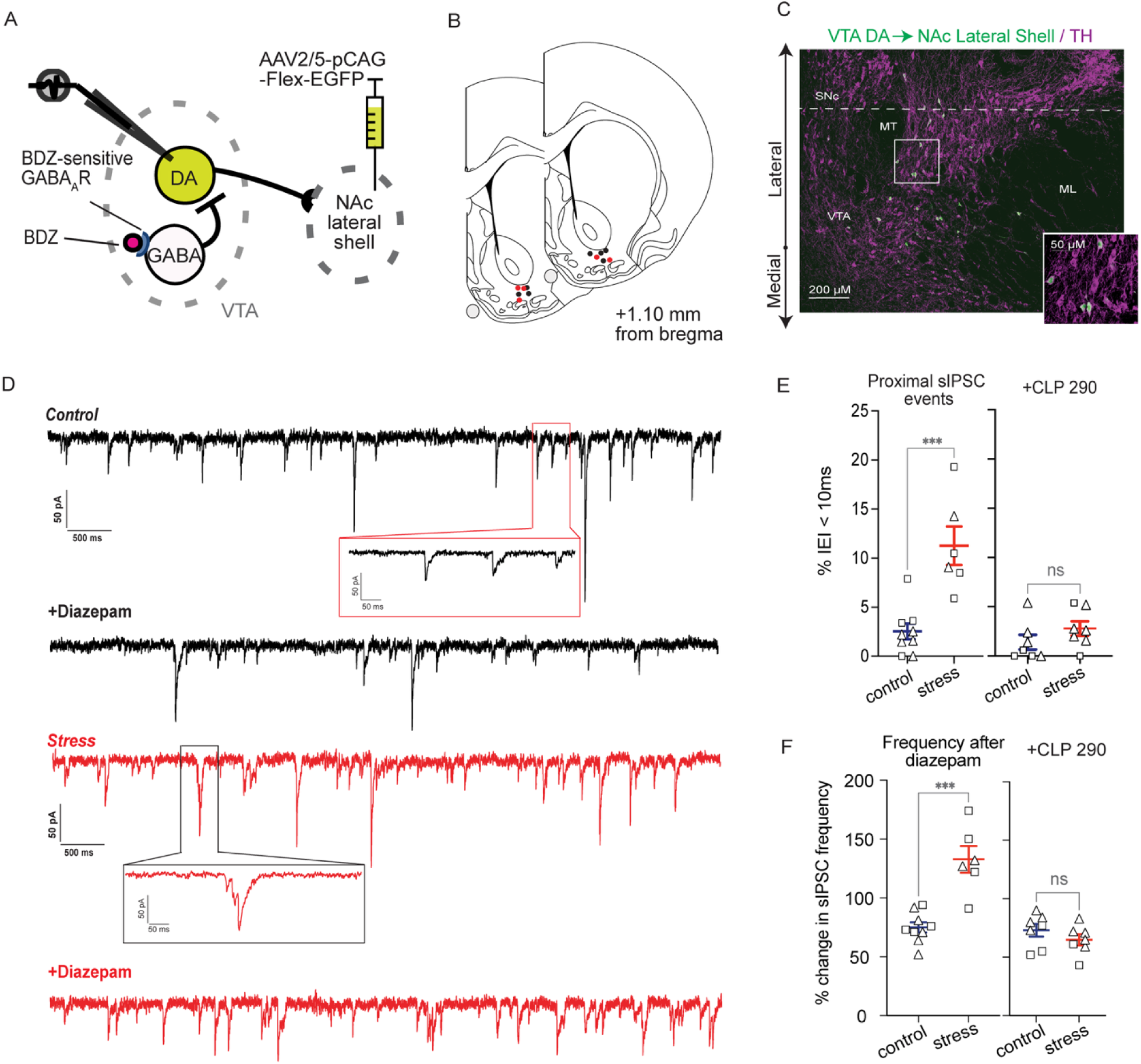
Stress-induced KCC2 downregulation alters GABAergic input onto NAc lateral shell-projecting DA neurons. A) TH-Cre rats received injections of a Cre-dependent GFP-expressing AAV into the NAc lateral shell to retrogradely label DA neurons. sIPSCs were recorded from GFP-labeled VTA DA neurons using whole-cell patch-clamp. Diazepam was applied to potentiate GABA_A_ receptor function in midbrain GABA neurons. B) AAV injection sites in the NAc lateral shell were comparable between control (black circles) and stressed (red circles) rats. C) Immunostaining showing that NAc lateral shell-projecting DA neurons (TH, magenta; eGFP, green) were primarily localized in the lateral VTA of TH-Cre rats. D) Representative sIPSC traces before and after diazepam in control (black) and stressed (red) rats. Insets show stress-induced shifts in inter-event interval distributions. E) Left: Lateral shell-projecting DA neurons from stressed animals exhibited a higher percentage of proximal sIPSCs (inter-event intervals <10 ms) compared to controls (***p = 0.0004, two-sided t-test, n = 9 cells/5 control rats, n = 6 cells/3 stressed rats). Right: Midbrain slice incubation with CLP290 prevented this effect (control: n = 7 cells/4 rats, stress: n = 7 cells/3 rats, p = 0.2071). Triangles represent females and squares represent males. F) Stress exposure significantly increased diazepam-induced sIPSC frequency in lateral shell-projecting DA neurons compared to controls (***p = 0.0001, two-sided t-test, n = 9 cells/5 control rats, n = 6 cells/3 stressed rats). CLP290 incubation prevented this effect (control: n = 7 cells/4 rats, stress: n = 7 cells/3 rats, p = 0.2810). Triangles represent females and squares represent males.

### 3.3 GABA input to NAc core and medial shell projecting DA neurons is not altered after stress

Given that DA signaling in the NAc core and medial shell can contribute to the acquisition of reward-related behaviors (Corre et al., 2018; Flagel et al., 2011; Ito et al., 2008), we next examined whether stress-induced KCC2 downregulation alters inhibitory transmission onto VTA DA neurons projecting to these regions. To do so, we retrogradely labeled DA neurons by injecting TH-Cre male and female rats with a Cre-dependent GFP-expressing AAV into the NAc core and medial shell (Figures 3A-C). Animals were then subjected to 1 hour of restraint stress, and sIPSCs were recorded from GFP-positive DA neurons approximately 15 hours later. In contrast to lateral shell-projecting neurons, DA projections to the NAc core and medial shell (Figures 3D, G) showed no significant stress-induced changes in the proportion of sIPSCs with interevent intervals <10 ms (Figures 3E, H): core projections: 3.9 ± 0.4% in stressed versus 3.3 ± 0.9% in control rats; and medial shell projections: 2.6 ± 0.7% in stressed versus 2.9 ± 1% in controls. Neither the total sIPSC frequency nor amplitude differed between stressed and control animals (core projections: 5.8 ± 1.5 Hz and 36.7 ± 3.6 pA after stress versus 5.6 ± 0.9 Hz and 40.3 ± 4.5 pA in control rats, p = 0.9396 and 0.5405, respectively; and medial shell projections: 3.1 ± 0.7 Hz and 53.2 ± 8.9 pA after stress versus 2.9 ± 0.8 Hz and 50.8 ± 6.0 pA in control rats, p = 0.8947 and 0.8227, respectively). Moreover, bath application of diazepam decreased sIPSC frequency similarly in both stressed and control animals (Figures 3F and 3I): core projections 70.9 ± 3.8% in stressed animals versus 76.3 ± 7.6% of basal frequency in control rats; and medial shell projections: 78 ± 5.2% in stressed animals versus 71.7 ± 4.9% of basal frequency in control rats. Finally, diazepam did not alter sIPSC amplitude in either stressed or control animals (core projections: 98.5 ± 12.1% of basal amplitude in stressed versus 90.7 ± 7.2% in controls, p = 0.5834; and medial shell projections: 94.4 ± 10.6% in stressed versus 89.3 ± 4.9% in control animals, p = 0.6412), indicating no stress-induced changes in pre- or postsynaptic GABAergic transmission. Together, these findings reveal a striking projection-specific effect of stress-induced KCC2 downregulation within mesolimbic circuits, selectively impacting VTA GABAergic regulation of DA neurons projecting to the NAc lateral shell, while sparing projections to the core and medial shell.

**Figure 3.**
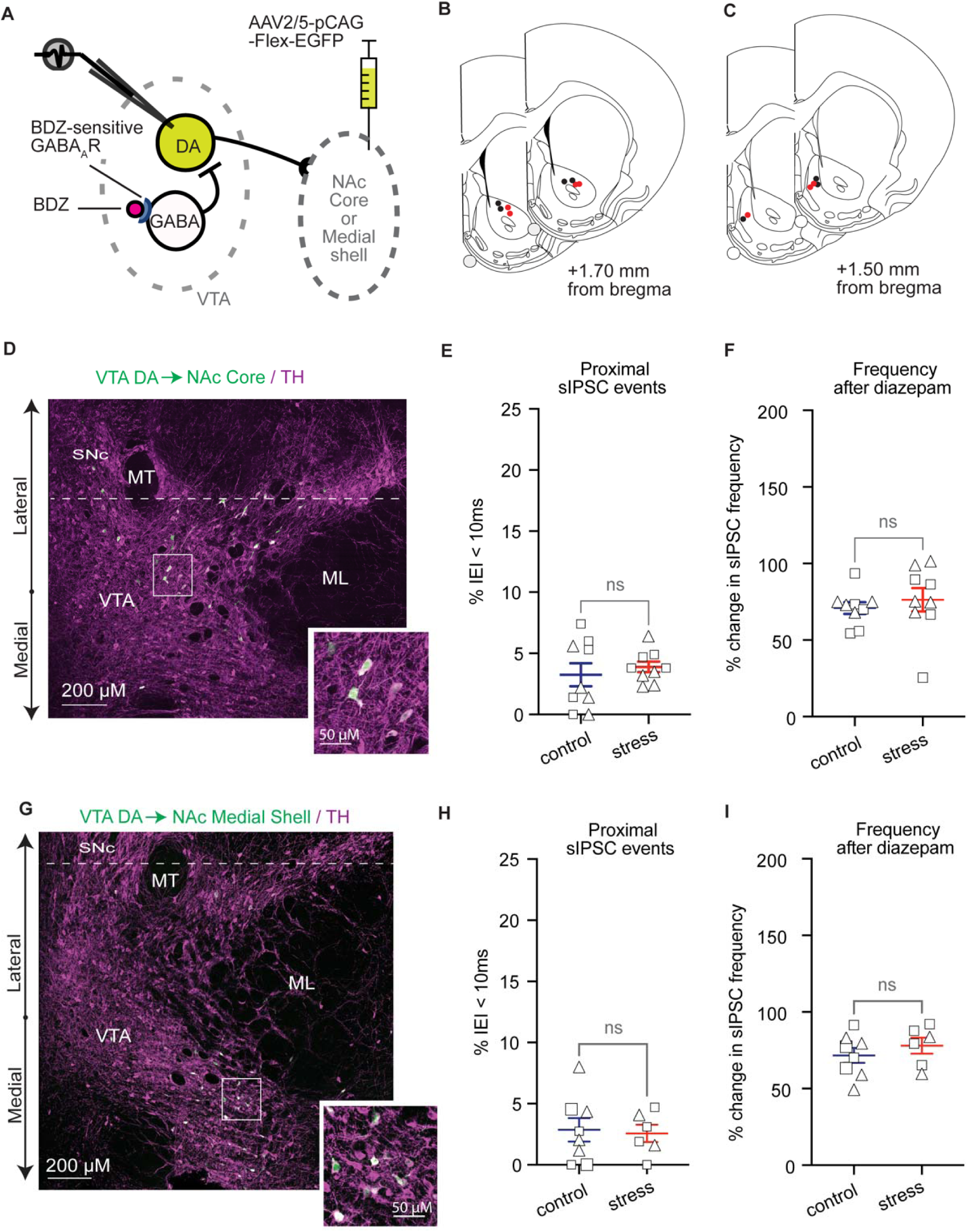
GABA input to NAc core- and medial shell-projecting DA neurons is not altered by stress. A) TH-Cre rats received injections of a Cre-dependent GFP-expressing AAV to retrogradely label DA neurons. sIPSCs were recorded from labeled VTA DA neurons using whole-cell patch-clamp. Diazepam was applied to potentiate GABA_A_ receptor function. B-C) Histological verification showing that AAV injection sites were comparable between control (black circles) and stressed (red circles) rats in the NAc core (B) and medial shell (C). D) Immunostaining showing eGFP (green) expression in VTA DA neurons (TH, magenta) following injection of Cre-dependent eGFP into the NAc core of TH-Cre rats. E-F) NAc core-projecting DA neurons from stressed animals did not differ from controls in the percentage of proximal sIPSCs (p = 0.54870), or in diazepam-induced sIPSC frequency (p = 0.5351; two-sided t-test; control: n = 9 cells/4 rats; stress: n = 9 cells/4 rats). Triangles represent females and squares represent males. G) Immunostaining showing eGFP (green) expression in VTA DA neurons (TH, magenta) following injection of Cre-dependent eGFP into the NAc medial shell of TH-Cre rats. H-I) Stress did not alter the percentage of proximal sIPSCs (p = 0.8207) or diazepam-induced sIPSC frequency (p = 0.4007) in medial shell-projecting DA neurons (two-sided t-test; control: n = 8 cells/4 rats; stress = 6 cells/4 rats). Triangles represent females and squares represent males.

### 3.4 Stress-induced KCC2 downregulation increases GABA input to VTA and SNc DA neurons projecting to medial, but not lateral sites of the dorsal striatum

Because DA signaling within the dorsomedial and dorsolateral striatum (DMS and DLS) contribute to reinforcement via action-outcome or habitual learning (Balleine and O’Doherty, 2010; Robinson et al., 2007a; Yin and Knowlton, 2006), we next examined whether stress and KCC2 downregulation alter GABAergic inputs to midbrain DA neurons projecting to these regions. Male and female TH-Cre rats received Cre-dependent GFP AAV injections in the DMS and DLS (Figures 4A-C), were subjected to 1-hour restraint stress, and sIPSCs were recorded in retrogradely labeled DA neurons approximately 15 hours later. In DMS-projecting DA neurons (Figure 4D), stress did not alter overall sIPSC frequency or amplitude: 4.2 ± 1.1 Hz and 26.8 ± 2.3 pA after stress versus 5.9 ± 1.2 Hz and 35.7 ± 4.1 pA in controls, p = 0.3381 and 0.0971, respectively. However, stress increased the proportion of temporally proximal events (Figure 4E, left; 18.7 ± 0.9% in stressed versus 7.9 ± 1.3% in controls) and potentiated diazepam-induced sIPSC frequency (Figure 4F, left; 128.4 ± 8.8% of baseline after stress versus 77.2 ± 5.7% in controls). No differences in sIPSC amplitude after diazepam were observed (90.4 ± 4.0% for stressed vs. 99.3 ± 6.9% for controls, p = 0.3048). Pre-incubation of midbrain slices with the KCC2 activator CLP290 (10 μM, 1 hour) normalized stress-induced changes, both in temporal proximity of sIPSCs (Figures 4E, right: 6 ± 0.7% in stressed versus 6.1 ± 0.7% in controls) and in diazepam-induced sIPSC potentiation (Figures 4F, right; 70.9 ± 6.7% in stressed versus 80.2 ± 7.0% in controls).

**Figure 4.**
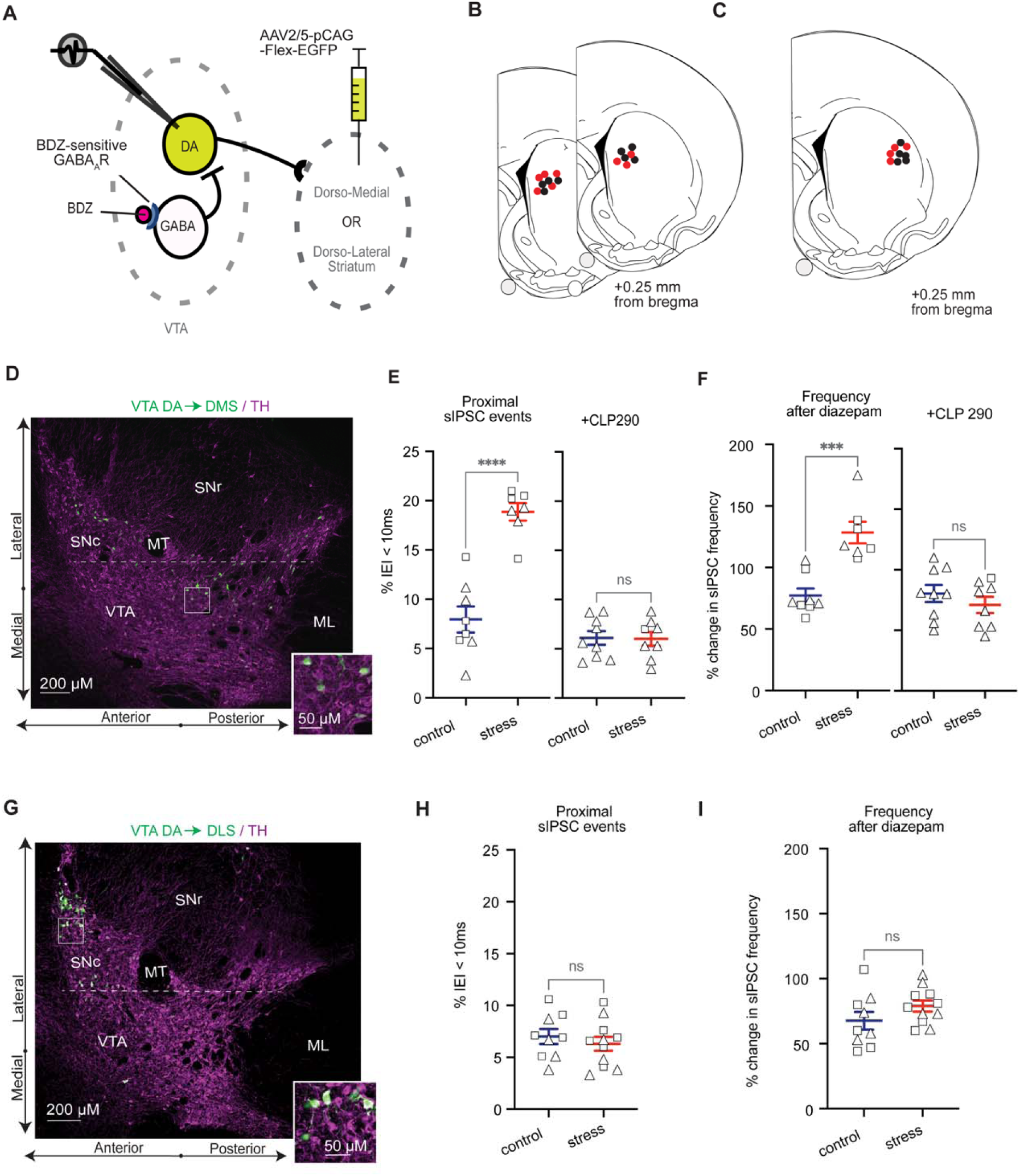
Stress-induced KCC2 downregulation alters GABA input onto DA neurons projecting to medial, but not lateral, dorsal striatum. A) TH-Cre rats were injected with a Cre-dependent AAV expressing GFP to retrogradely label DA neurons projecting to the dorsomedial (DMS) or dorsolateral striatum (DLS). sIPSCs were recorded from labeled VTA DA neurons using whole-cell patch-clamp. Diazepam was applied to enhance GABA_A_ receptors in VTA GABA neurons. B-C) Injection sites in the DMS (B) and DLS (C) were comparable between control (black circles) and stressed (red circles) animals. D) Immunostaining showing that DMS-targeted injections resulted in eGFP (green) expression in DA neurons (TH, magenta) located in the lateral VTA and medial SNc. E) Left: DMS-projecting DA neurons from stressed animals (n = 7 cells/5 rats) exhibited a higher proportion of proximal sIPSCs (inter-event interval < 10 ms) compared to controls (n = 8 cells/4 rats; ****p < 0.0001, two-sided t-test). Right: Incubation of midbrain slices with CLP290 prevented this effect (p = 0.9251; two-sided t-test; control: n = 9 cells/4 rats; stress: n = 8 cells/3 rats). Triangles represent females and squares represent males. F) Stress exposure significantly increased diazepam-induced sIPSC frequency in DMS projecting DA neurons compared to control, non-stressed animals (***p = 0.0002; two-sided t-test; control: n = 8 cells/4 rats; stress = 7 cells/5 rats). CLP290 incubation prevented this effect (p = 0.3537; two-sided t-test; control = 9 cells/4 rats); stress = 8 cells/3 rats). Triangles represent females and squares represent males. G) Immunostaining showing eGFP (green) expression in SNc DA neurons (TH, magenta) following DLS-targeted injections. H-I) In DLS-projecting DA neurons, stress did not significantly alter the proportion of proximal sIPSCs (p = 0.4865) or diazepam-induced sIPSC frequency (p = 0.1599; two-sided t-tests; control: n = 9 cells/5 rats); stress: n = 11 cells/4 rats). Triangles represent females and squares represent males.

In contrast, DLS-projecting DA neurons (Figure 4G) were unaffected by acute stress. There were no changes in the proportion of proximal sIPSCs (6.3 ± 0.7% in stressed versus 7.0 ± 0.7% in controls; Figure 4H) or in diazepam-induced attenuation of sIPSC frequency (78.9 ± 4.2% in stressed vs. 67.6 ± 6.9% in controls, n = 9, 11 cells in 5, 4 rats; Figure 4I). Likewise, overall sIPSC frequency and amplitude were unchanged (3.1 ± 0.3 Hz and 36.1 ± 3.0 pA in stressed vs. 2.4 ± 0.4 Hz and 29.8 ± 2.2 pA in controls, p = 2.400 and 0.1181, respectively). These results indicate that, within the VTA/SNc, stress induces a pathway-specific effect of KCC2 downregulation, selectively altering GABAergic inputs to DMS-projecting DA neurons.

### 3.5 KCC2 activation prevents stress-induced potentiation of self-administration acquisition

Previous work demonstrated that KCC2 downregulation impairs inhibitory signaling onto DA neurons, enhancing DA firing and release (Woo et al., 2025). Our new findings (Figures 2-4) show that stress-induced KCC2 downregulation selectively alters mesolimbic and nigrostriatal DA pathways involved in cue-reward and action-outcome learning, both of which may contribute to enhanced acquisition of sucrose self-administration (Figure 1). We therefore hypothesized that upregulating KCC2 with the activator CLP290 could prevent stress-induced facilitation of sucrose self-administration. To test this, male and female rats pre-exposed to a 1-hour restraint stress received intraperitoneal injections of CLP290 (10 mg/kg) or vehicle (20% β-cyclodextrin) approximately 1 hour before the self-administration task on alternating days during the first 6 days of training (Figure 5A). Compared to stressed rats receiving vehicle (Figure 5B, red), CLP290-treated stressed rats showed significantly reduced sucrose self-administration (Figure 5B, blue). These data were indistinguishable from the non-stressed control group (Figure 5B, dotted black line). Inactive lever responding did not differ between groups (Figure 5C). Accordingly, the difference between active and inactive lever presses was significantly lower in CLP290-treated stressed animals compared to vehicle-treated controls (Figure 5D, 21 ± 6.2 in vehicle-administered versus 6.3 ± 3.1 in CLP290-administered stressed rats). Using the same acquisition criterion as in Figure 1 (≥10 presses on the active versus inactive lever for at least two consecutive days), 63.6% of vehicle-treated stressed animals acquired sucrose self-administration within the first six sessions, compared to 28.6% of CLP290-treated stressed animals (Figure 5D). Among those meeting this criterion, CLP290- and vehicle-treated stressed rats exhibited similar responding during days 7-10 (Figures 5E-G).

**Figure 5.**
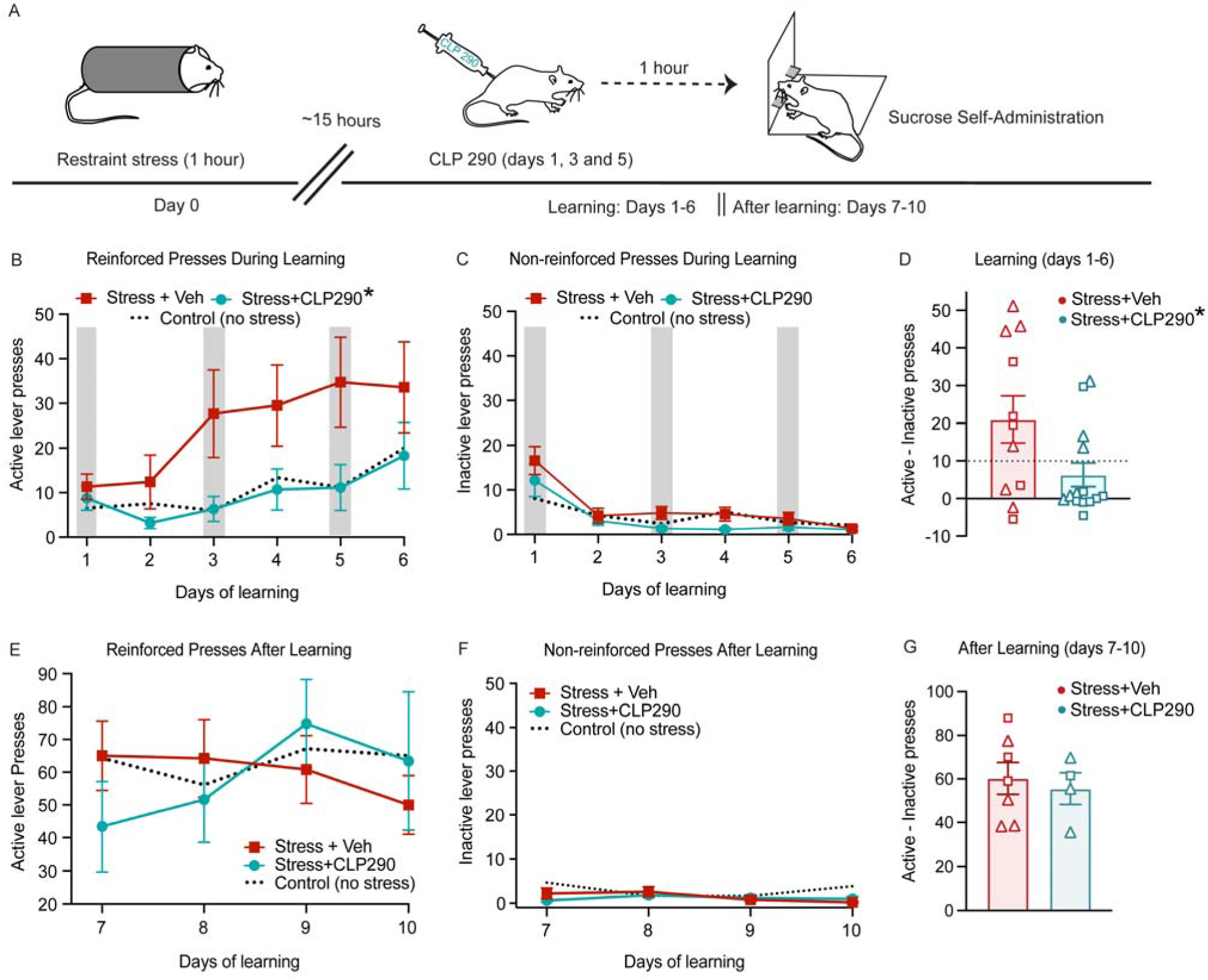
Stress-induced potentiation of acquisition of self-administration behavior is prevented by CLP290. A) Stressed rats received systemic CLP290 or vehicle 1 h before self-administration sessions 1, 3, and 5. Active and inactive lever presses were measured during the first 6 days of training and for an additional 4 days in animals that acquired sucrose self-administration. B) CLP290-treated stressed rats showed significantly reduced active lever pressing compared with vehicle-treated stressed rats (*p = 0.0250, RM ANOVA, treatment: F (1, 23) = 5.753, n = 14 CLP290 treated, and 11 vehicle-treated stressed rats). Active lever pressing in non-stressed control rats is shown for comparison (dotted line). C) Inactive lever presses did not differ between CLP290 and vehicle-treated stressed rats (p = 0.1095, RM ANOVA). D) The mean daily difference between active and inactive lever presses was lower in CLP290-treated stressed rats compared with vehicle-treated stressed rats (*p = 0.0344, two-sided t-test, n = 11, 14 rats). Triangles represent females and squares represent males. The dashed line indicates the acquisition criterion (≥10 presses on the active versus inactive lever). E-F) In stressed rats that acquired sucrose self-administration within the first 6 sessions, active (E) and inactive (F) lever pressing did not differ between treatment groups (E: p = 0.7828; F: p = 0.6454; RM ANOVA, n = 5, 5 rats). G) The mean daily difference between active and inactive lever presses on days 7-10 did not differ between CLP290- and vehicle-treated stressed rats (p = 0.6844, two-sided t-test). Triangles represent females and squares represent males.

## 4. Discussion

In the present study, we show that acute stress selectively reshapes GABAergic input onto distinct DA circuits to facilitate the acquisition of reward-seeking behavior. Specifically, acute restraint stress increased active lever pressing during the early stages of sucrose self-administration and altered GABAergic inputs onto DA neurons projecting to the NAc lateral shell and DMS, while not affecting projections to the DLS and other accumbal subregions. These circuit-specific effects were associated with downregulation of KCC2 in GABAergic inputs. Importantly, pharmacological enhancement of KCC2 function prevented the stress-induced facilitation of sucrose self-administration, supporting KCC2-dependent modulation of inhibitory input onto DA neurons as a key mechanism by which stress biases reward-related learning.

We demonstrate that pre-exposure to acute restraint stress potentiates the acquisition of operant sucrose self-administration. Animal studies generally demonstrate that acute stress enhances reward learning, although these effects depend on multiple factors, including the type and timing of the stressor and the behavioral paradigm employed (Schettino et al., 2024). In our study, a 1-hour restraint was administered approximately 15 hours before the sucrose self-administration task, allowing us to assess the lasting neural circuit consequences of stress, rather than its immediate effects. Importantly, we found that stress selectively enhanced the early stages of behavior acquisition (more animals acquired the operant behavior after stress) but once the behavior was established, their operant pressing was not different from that of control animals. This finding aligns with studies on associative learning, where stress has been shown to enhance Pavlovian conditioning to rewarding stimuli (Stelly et al., 2020). In the context of operant conditioning, stress exposure has been shown to potentiate the acquisition of addictive drug self-administration behaviors (e.g., cocaine, morphine, ethanol) (Carter et al., 2020; Garcia-Keller et al., 2016; Ostroumov et al., 2016). With regard to non-drug rewards, human studies suggest that pre-task stress exposure can enhance reinforcement learning; however, findings have been considerably more variable than those reported in animal studies (Lighthall et al., 2013; Schettino et al., 2024). Yet, to our knowledge, this is the first study showing potentiating effect of acute stress on the acquisition phase of non-drug self-administration in rodents.

Our findings suggest that stress enhances the acquisition of sucrose self-administration through KCC2 downregulation in GABAergic inputs onto DA neurons. In previous studies, we demonstrated that stress downregulates KCC2 in VTA GABA neurons, which provide a major inhibitory input to DA neurons (Doyon et al., 2013; Ostroumov et al., 2016). Stress-induced KCC2 downregulation has been shown to increase the consumption of addictive drugs, particularly during the early stages of self-administration (Ostroumov et al., 2016). More recently, we demonstrated that KCC2 downregulation is necessary for the acquisition of Pavlovian learning (Woo et al., 2025). Critically, the potentiation of learning involves the synchronization of GABAergic input onto DA neurons, an effect also observed in the current study. This synchronization of VTA GABA neurons results in increased phasic firing of DA neurons, which is necessary for the acquisition of reward-related behaviors (Woo et al., 2025). Based on these findings and prior work, we hypothesize that stress-induced synchronization of GABAergic input onto DA neurons (Figures 2 and 4) potentiates reward-related learning by enhancing behaviorally relevant DA signaling through KCC2 downregulation in local midbrain GABA neurons.

An important consideration is that, in addition to local inhibition, DA neurons also receive GABAergic input from other brain regions. However, the observed effect of CLP290 in reversing stress-induced changes in sIPSCs excludes the contribution of distal GABAergic inputs to these changes. Our midbrain slice preparation preserves local GABA neurons but only contains neuronal terminals from distal GABAergic inputs. Since KCC2 is expressed in dendrites and somata, but not in terminals, CLP290 exposure would enhance KCC2 function specifically in local GABA neurons, with no effect on terminals. Thus, the reversal of stress-induced changes in sIPSCs by CLP290 (Figures 2 and 4) suggests that stress downregulates KCC2 specifically in midbrain GABA neurons, which provide the major GABAergic input to DA neurons, including neurons in the VTA, substantia nigra pars reticulata, and rostromedial tegmental nucleus. Our previous findings show stress-induced KCC2 downregulation in VTA GABA neurons, but future studies could explore whether acute stress similarly impairs KCC2 function in other midbrain GABAergic populations.

Our findings demonstrate notable circuit specificity within the mesolimbic and nigrostriatal systems, wherein stress-dependent KCC2 downregulation preferentially impacts inhibitory transmission onto DA neurons projecting to the NAc lateral shell and DMS. These pathways are associated with partially distinct aspects of reward-related behavior. Lateral mesolimbic projections, including those to the NAc lateral shell, have been implicated in associative learning, reinforcement, and incentive salience attribution, particularly in linking environmental cues to rewarding outcomes (Chen et al., 2023; de Jong et al., 2024; Smedley et al., 2019; Yang et al., 2018). In contrast, medial nigrostriatal DA projections and the DMS are critically involved in goal-directed action control, including action-outcome contingency learning and the selection of actions based on their consequences (McCane et al., 2021; Parker et al., 2016; Robinson et al., 2007b; Yin et al., 2005). Accordingly, stress-induced KCC2 downregulation in GABAergic inputs onto NAc lateral shell- and DMS-projecting DA neurons may facilitate acquisition of operant sucrose self-administration by enhancing the association of environmental cues and/or actions with reward. However, future studies are required to establish direct causal links between midbrain KCC2 downregulation, DA signaling changes in projection-defined pathways, and operant learning behavior.

Another finding that warrants further investigation is the absence of stress-dependent KCC2 downregulation effects in NAc medial shell-, core-, and DLS-projecting DA neurons. This lack of effect may reflect differences in the organization of inhibitory inputs onto distinct DA neuron subpopulations (Soden et al., 2020). For example, previous work has suggested that NAc lateral shell-projecting DA neurons receive stronger local GABA_A_ receptor-mediated input compared to more medial accumbal projections (Yang et al., 2018). Accordingly, the absence of stress effects in these DA neuron populations may arise from differences in the strength, density, or functional efficacy of GABAergic synaptic inputs. Furthermore, unaffected DA neurons may be preferentially innervated by distinct subclasses of midbrain GABA neurons that differ in KCC2 expression or in their sensitivity to stress-related modulation (Bouarab et al., 2019).

In summary, the present findings indicate that stress enhances the acquisition of operant reward self-administration and reveal that stress-induced inhibitory plasticity differentially affects mesolimbic and nigrostriatal DA projections. These results extend prior work showing that acute stress enhances associative classical conditioning (Stelly et al., 2020) by demonstrating that stress also facilitates instrumental learning of actions that produce reward. Our findings are also consistent with heterogeneity across mesolimbic and nigrostriatal pathways, in which projection-defined DA neuron populations within the VTA and SNc exhibit distinct circuit organization and regulatory inputs. The observation that these projection-defined populations are differentially modulated by stress-induced changes in GABAergic input further supports the idea that these anatomical pathways may differentially contribute to distinct components of reward processing. Reward learning is often dysregulated in neuropsychiatric disorders, including stress-related disorders and addiction. Therefore, elucidating the molecular mechanisms by which stress facilitates reward learning may identify potential therapeutic targets for these conditions.

## Data availability statement

All data published in this article is available upon request.

## CRediT authorship contribution statement

**Helena C. Schuch:** Conceptualization, Methodology, Investigation, Data curation, Formal Analysis, Writing – original draft, Writing – Review & Editing. **Joyce Woo:** Investigation, Formal Analysis, Data curation, Writing – Review & Editing. **Hannah Kugler:** Investigation, Data curation. **Kelly Jiang:** Investigation, Data curation. **Alexey Ostroumov:** Conceptualization, Formal Analysis, Supervision, Project administration, Funding acquisition, Writing - Review & Editing.

## Acknowledgements

We thank Anna Pearson for helpful discussions and thoughtful feedback on the manuscript.

